# Inter- and intra-subject similarity in network functional connectivity across a full narrative movie

**DOI:** 10.1101/2024.05.14.594107

**Authors:** Lisa N. Mochalski, Patrick Friedrich, Xuan Li, Jean-Philippe Kröll, Simon B. Eickhoff, Susanne Weis

## Abstract

Naturalistic paradigms, such as watching movies during functional magnetic resonance imaging (fMRI), are thought to prompt the emotional and cognitive processes typically elicited in real life situations. Therefore, naturalistic viewing (NV) holds great potential for studying individual differences. However, in how far NV elicits similarity within and between subjects on a network level, particularly depending on emotions portrayed in movies, is currently unknown.

We used the studyforrest dataset to investigate the inter– and intra-subject similarity in network functional connectivity (NFC) of 14 meta-analytically defined networks across a full narrative, audio-visual movie split into 8 consecutive movie segments. We characterized the movie segments by valence and arousal portrayed within the sequences, before utilizing a linear mixed model to analyze which factors explain inter– and intra-subject similarity.

Our results showed that the model best explaining inter-subject similarity comprised network, movie segment, valence and a movie segment by valence interaction. Intra-subject similarity was influenced significantly by the same factors and an additional three-way interaction between movie segment, valence and arousal.

Overall, inter– and intra-subject similarity in NFC were sensitive to the ongoing narrative and emotions in the movie. Lowest similarity both within and between subjects was seen in the emotional regulation network and networks associated with long-term memory processing, which might be explained by specific features and content of the movie. We conclude that detailed characterization of movie features is crucial for NV research.

## 1. Introduction

Understanding how individual differences in brain architecture shape personality, cognitive abilities and socio-affective traits is a constant endeavor in cognitive neuroscience. The growing interest in individual differences research has led to the development of new paradigms that may allow for novel insights into individual brain architecture. Naturalistic viewing (NV) is a promising tool for designing more ecologically valid fMRI studies, thus providing the opportunity to measure individual differences under beneficial circumstances (Vanderwal et al., 2017). However, the potential and limitations of naturalistic viewing for the study of individual differences are not yet fully understood.

Past fMRI research mainly followed one of two approaches to uncover the brain’s functional architecture, each of which comes with their own strengths and constraints. Traditionally, task-based fMRI has been essential for studying the neural correlates of cognition based on paradigms designed to target specific cognitive processes in a highly controlled environment (Dosenbach et al., 2006). However, the highly controlled nature of task fMRI leads to rather artificial participant experiences and thus sacrifices ecological validity for precision (Kingstone, Smilek, Eastwood, 2008; Delgado-Herrera, Reyes-Aguilar, Giordano, 2021; Reggente et al., 2018; van Atteveldt et al., 2018). Alternatively, the intrinsic functional architecture of the brain has been studied using resting state (RS) fMRI, in which participants are scanned without any task or external stimulus. However, in this paradigm, subjects are left to follow their own thoughts, which impacts brain states in a way that is difficult to control (Gonzalez-Castillo, Kam, Hoy & Bandettini, 2021).

While both task and RS-fMRI possess their strengths, their constraints have led to the development of NV paradigms as an alternative for studying individual differences in neuroimaging research. In NV, subjects are presented with movies or movie segments while undergoing MRI scanning, without any additional task instructions beyond attending to the movie. Movies are assumed to induce brain states similar to those evoked in real life situations, because movies are likewise complex, dynamic and continuous. NV thus aims to deliver ecologically valid stimulation to the brain as far as this is possible within the confines of a MR scanner. Compared to task-based fMRI, participants’ brain states are less constrained while experiencing a less artificial situation during NV (Finn et al., 2017). In combination with advantages of participant engagement and compliance, this additionally creates interest in movie fMRI for clinical applications (Eickhoff, Milham & Vanderwal, 2020) or specific populations such as children (Vanderwal et al., 2019). Simultaneously, movies offer rich, multi-modal stimulation that induces brain states in a more constrained way than RS-fMRI does.

Therefore, movies allow for employing the whole range of analyses that are typical for task fMRI and RS-fMRI (Spiers & Maguire, 2007; Vanderwal et al., 2019; Nastase et al., 2019). NV can induce high inter– and intra-subject correlations in activity time courses of various cortices (Hasson et al., 2004), which are dependent on features and content of the movie stimulus (Hasson et al., 2008; Lerner, Honey, Silbert & Hasson, 2011). These inter– and intra-subject correlations in brain activity are affected by the narrative coherence of a movie stimulus, as backward presentation of a movie decreases these correlations (Hasson, Malach & Heeger, 2009). Moreover, movie stimuli can be edited to influence similarity as shown by higher inter-subject correlations in professionally produced movies than unedited, real-life movies (Hasson, Malach & Heeger, 2009). A direct comparison between different movie stimuli and RS indicated that a complex movie with social interactions yielded higher inter-subject correlations compared to an abstract, non-verbal movie, which in turn led to higher inter-subject correlations than RS (Vanderwal, Kelly, Eilbott, Mayes, & Castellanos, 2015).

Moreover, movies are effective in eliciting emotions, including complex and differentiated emotional states (Gross & Levenson, 1995; Westermann et al., 1996; Schaefer et al., 2010; Adolphs et al., 2016). Emotions might be induced using various features in the movie, such as facial expressions, gestures, body postures, speech characteristics or context cues (Skerry and Saxe, 2014). In movies with social content, the emotions portrayed by characters in the movie are important cues for eliciting emotions in viewers (Labs et al., 2019, Lettieri et al., 2019), making portrayed emotions an important stimulus feature in NV studies.

According to Finn and colleagues (2017), a major argument why movie fMRI might be an excellent paradigm for studying individual differences is the assumed beneficial ratio of inter-to intra-subject correlation induced by movies: On the one hand movies represent a common cognitive reference frame for subjects’ brain states, thus decreasing irrelevant inter-subject variability, while on the other hand retaining a subject’s most identifying features. This is denoted by low intra-subject variability, i.e. subjects being similar to themselves, for example, over the course of watching a movie or when watching the same movie in two separate sessions (Finn et al., 2017). Concordantly, a study by Vanderwal and colleagues (2017) showed that movies overall significantly decreased both inter– and intra-subject variability on a whole-brain level compared to RS, thus lending support to the idea that NV might preserve or even enhance individual differences in functional connectivity (Vanderwal, et al. 2017). A recent study found that influences of NV on inter– and intra-subject similarity in NFC is dependent on the brain network and stimulus (Kröll et al., 2023), however, not all factors contributing to the effects of movies on inter– and intra-subject similarity of NFC have been investigated. For example, the impact of specific content, such as emotions portrayed in movies, is still unknown.

In contemporary neuroscience, functional networks consisting of distributed but interacting brain regions are often viewed as the foundation of cognition functions (Eickhoff and Grefkes, 2011), thus allowing for an interpretation of interactions between various brain regions with respect to specific cognitive domains. Given that movies are complex and multimodal stimuli that elicit widely distributed brain activity, a network perspective might help to untangle the effect of movies on specific cognitive functions. Specifically, studying the functional connectivity between the brain regions constituting these networks (i.e. network functional connectivity, NFC) might grant insights into the effects of NV on brain function. While there are various methods to define functional networks (e.g. Fox et al., 2005; Power et al., 2011; Smith et al., 2009; Yeo et al., 2011; Schaefer et al., 2018, Pervaiz et al., 2020), one approach that instrumentalizes the body of existing knowledge about specific cognitive processes is the use of meta-analysis. Coordinate-based meta-analyses of neuroimaging data (e.g. activation likelihood estimation; Eickhoff et al., 2009) identify brain locations that are consistently activated during cognitive tasks across various studies. Converging results from many studies using different tasks to study the same cognitive function leads to a robust mapping of function-related brain coordinates (Eickhoff et al., 2012). In turn, reliably co-activated regions can be assumed to constitute a network that is engaged with the specific cognitive function (Fox et al., 2015). Various meta-analytical networks have been characterized, covering different psychological domains, and have been proven useful for gaining insight into the role of brain regions in a network perspective (Igelström & Graziano, 2017), robustly assessing the neural basis of cognitive functions (Gross, 2015; Etkin, Büchel & Gross, 2015; Binder & Desai, 2011, Margulies et al., 2016) and therefore laying the ground for further experimental work (Morawetz et al., 2017). Studies using meta-analytical networks to predict personality scores (Nostro et al., 2018) or classify participants according to their mental health status and age (Pläschke et al., 2017) yielded better or at least similar results to using whole-brain connectivity (Nostro et al., 2018), while improving interpretability. Therefore, meta-analytical networks provide an excellent basis for studying individual variability in different cognitive systems.

With respect to movie fMRI, Vanderwal and colleagues (2017) showed that the effect of movies is differentially distributed across the brain, with lower inter-than intra-subject variability in unimodal regions and higher inter-than intra-subject variability in heteromodal regions. However, a concrete comparison of these variabilities on the level of network functional connectivity (NFC) has rarely been done (but see Kröll et al., 2023). Furthermore, it is yet unclear which features of a movie stimulus influence inter– and intra-subject similarity. Emotions might play an important role in affecting similarity within and between subjects and might be best studied in a full narrative movie, which combines a long, overarching narrative and numerous distinct, individual scenes. The changing content and wide range of emotional scenes present within this narrative warrants a more detailed characterisation of the stimulus and its segments.

To fill this gap, the present study investigates inter– and intra-subject similarity in NFC over the course of a full narrative movie. In a first step, we compared different segments of the movie stimulus with regard to their portrayed valence and arousal, evincing differences in emotional content. Using linear mixed models (LMM), we analyzed how different factors, such as the narrative of the movie and portrayed valence and arousal, affect inter– and intra-subject similarity in NFC across 14 meta-analytically defined networks.

We expected to see differences in inter– and intra-subject similarity that are network-dependent, change over the course of the full narrative movie and are influenced by valence and arousal portrayed within the movie.

Delineating the factors influencing inter– and intra-subject similarity is an essential next step in NV research. While the number of available datasets is steadily increasing, there is still no systematic framework guiding the choice of stimuli due to a lack of studies uncovering the influence of specific stimulus features on brain measures (Grall & Finn, 2022). Using movie watching precisely and effectively necessitates a better understanding of which stimuli are best suited for elucidating which research questions (Eickhoff, Milham & Vanderwal, 2020). This study sheds light on the effects on inter– and intra-subject similarity in NFC of a movie stimulus that is unique in its length and overarching narrative.

## 2. Methods

### 2.1. Sample

This sample consisted of 15 native German-speaking participants (6 females, range 21-39 years) (Hanke et al., 2016). One subject, which we excluded, was an outlier in the intra-subject correlation analysis, leading to a sample size of 14 (6 females, age range 21-39 years. Please note that mean age cannot be reported, because only age ranges were reported for each participant). The Ethics committee of Otto-Von-Guericke University, Germany approved acquisition of the data in the “studyforrest” project. For a more detailed description of the sample, see Sengupta et al. (2016). The full dataset can be found under: https://github.com/psychoinformatics-de/studyforrest-data-phase2. A list of subjects can be found in the supplementary table S1.

### 2.2 MRI Data & preprocessing

fMRI data acquisition took place in a single session which included a short break in the middle. To keep the stimulus at a length of two hours, some scenes were cut. The movie stimulus represents a full narrative movie, as only scenes that were less relevant to the plot were cut, thus preserving the overarching story. For the purpose of data acquisition, the movie stimulus was separated into 8 segments of approximately 15 minutes each, taking scene boundaries into consideration. This lead to an unequal number of volumes acquired per segment, which were 451, 441, 438, 488, 462, 439, 542, and 338 for segments 1 – 8 respectively (see Hanke et al., 2014 for details and code on movie segment creation). The movie segments were shown in chronological order. For each segment, T2*-weighted echo-planar images (gradient-echo, 2 s repetition time (TR), 30 ms echo time, 90° flip angle, 1943 Hz/Px bandwidth, parallel acquisition with sensitivity encoding (SENSE) reduction factor 2, 35 axial slices, 3.0mm slice thickness, 80 x 80 voxels (3.0 x 3.0mm) in-plane resolution, 240 x 240 mm field-of-view, anterior-to-posterior phase encoding direction in ascending order, 10% inter-slice gap, whole-brain coverage) were acquired using a whole-body 3 Tesla Philips Achieva dStream MRI scanner equipped with a 32 channel head coil.

All downloaded data were minimally preprocessed as described in Hanke et al. (2016). In short, preprocessing steps included defacing, motion correction, reslicing and data interpolation using in-house codes that utilize the FSL toolkit. All codes are openly available under: https://github.com/psychoinformatics-de/studyforrest-data-aligned/tree/master/code. For precise information about observed motion and data quality analyses see Hanke et al. (2016). For this study, the native fMRI data were brought into MNI space using FSL’s applywarp function for subsequent NFC extraction.

### 2.3 Valence and Arousal measures

To characterize the movie segments with regards to the portrayed valence and arousal, we used the openly available data from Labs et al. (2015). This dataset contains annotations of portrayed emotions in the “Forrest Gump” movie stimulus used in the “studyforrest” dataset. A group of observers (n=9, German female university students) were asked to evaluate scenes of the movie in terms of valence (“positive” or “negative”) and arousal (“high” or “low”) portrayed by the movie characters. All scenes were presented in random order to allow observers to focus on current indicators of portrayed emotions without being influenced by, for example, the conveyed mood of the movie plot. To evaluate the consistency of evaluations between observers, Labs and colleagues calculated the inter-observer agreement (IOA). The IOA value describes the portion of observers indicating the presence of a specific attribute in a scene (Labs et al, 2015). As arousal and valence were measured on a bipolar scale (“positive” of “negative” valende, “low” or “high” arousal present), the timeseries of these attributes were calculated as the difference between the IOA timeseries of both expressions. That is, the IOA timeseries of arousal was calculated by subtracting the IOA timeseries of low arousal segments from the IOA timeseries of high arousal segments (Labs et al., 2015). The IOA is expressed as a value between 1 and –1, with “1” indicating perfect observer agreement regarding high arousal (or positive valence, respectively) and “-1” indicating perfect observer agreement regarding low arousal (or negative valence, respectively). IOAs were reported as a time series of the movie in correct order downsampled to 2 seconds, corresponding to the sampling rate of the fMRI data. We used code published by Lettieri et al. 2019 (https://osf.io/tzpdf/) to divide the IOA time series according to 8 movie segments for subsequent analyses.

### 2.4 Inter– and intra-subject similarity in functional networks

To investigate effects on a network level, we used 14 networks defined as sets of peak coordinates in different meta-analyses. These included the autobiographical memory (AM) network (Spreng, Mar & Kim, 2008), cognitive attention control (CogAC) network (Cieslik et al., 2015), extended multiple demand network (eMDN) (Camilleri et al., 2018), emotional scene and face processing (EmoSF) network (Sabatinelli et al., 2011), empathy network (Bzdok et al., 2012), theory of mind (ToM) network (Bzdok et al., 2012), emotion regulation (ER) network (Buhle et al., 2014), extended socio-affective default network (eSAD) (Amft et al., 2015), mirror neuron system (MNS) network (Caspers et al., 2010), motor network (Witt, Meyerand, & Laird, 2008), reward (Rew) network (Liu et al., 2011), semantic memory (SM) network (Binder et al., 2009), vigilant attention (VigAtt) network (Langner & Eickhoff, 2013), and the working memory (WM) network (Rottschy et al., 2012). A more detailed description of these networks are reported in the supplements (supplementary material S2). For each meta-analytical network, nodes were created by placing 6mm spheres around the peak coordinates (see supplementary material S3 for an overview of the peak coordinates and S4 for a figure of all nodes of all networks). The functional connectome of a given network was created using in-house MATLAB R2017a (The Mathworks Inc., 2017) code which computes the pairwise Pearson correlation between all nodes for each segment and each participant. This resulted in 1680 functional network connectomes (15 participants x 8 segments x 14 networks) saved as N-by-N matrices with N being the number of nodes.

To keep in line with previous studies (Vanderwal et al., 2015; Vanderwal et al., 2017; Finn et al., 2017; Nastase et al., 2019), we operationalized the inter– and intra-subject similarity as the Pearson correlation coefficients between functional connectomes within and between subjects. Inter– and intra-subject similarity were computed per network, segment and subject as depicted in Figure 1. All computations are based on the unique connections between nodes (i.e. the lower triangle of the NFC matrix) and exclude all auto-correlations. For inter-subject similarity, we first computed the correlations between the NFC of one subject and all other subjects. After Fisher Z-transformation of the correlation coefficient, they were averaged and re-transformed, resulting in one value representing inter-subject similarity for the respective subject in the given segment and network. For calculating intra-subject similarity of a given subject and segment, we computed the correlations between NFCs of this segment and every other segment of the subject. The correlation values were Fisher’s z-transformed, averaged, and reverted to r-values, resulting in one value representing intra-subject similarity for the respective subject in the given segment and network. Both inter– and intra-subject similarity were calculated based on Pearson correlation coefficients.

**Figure 1:**
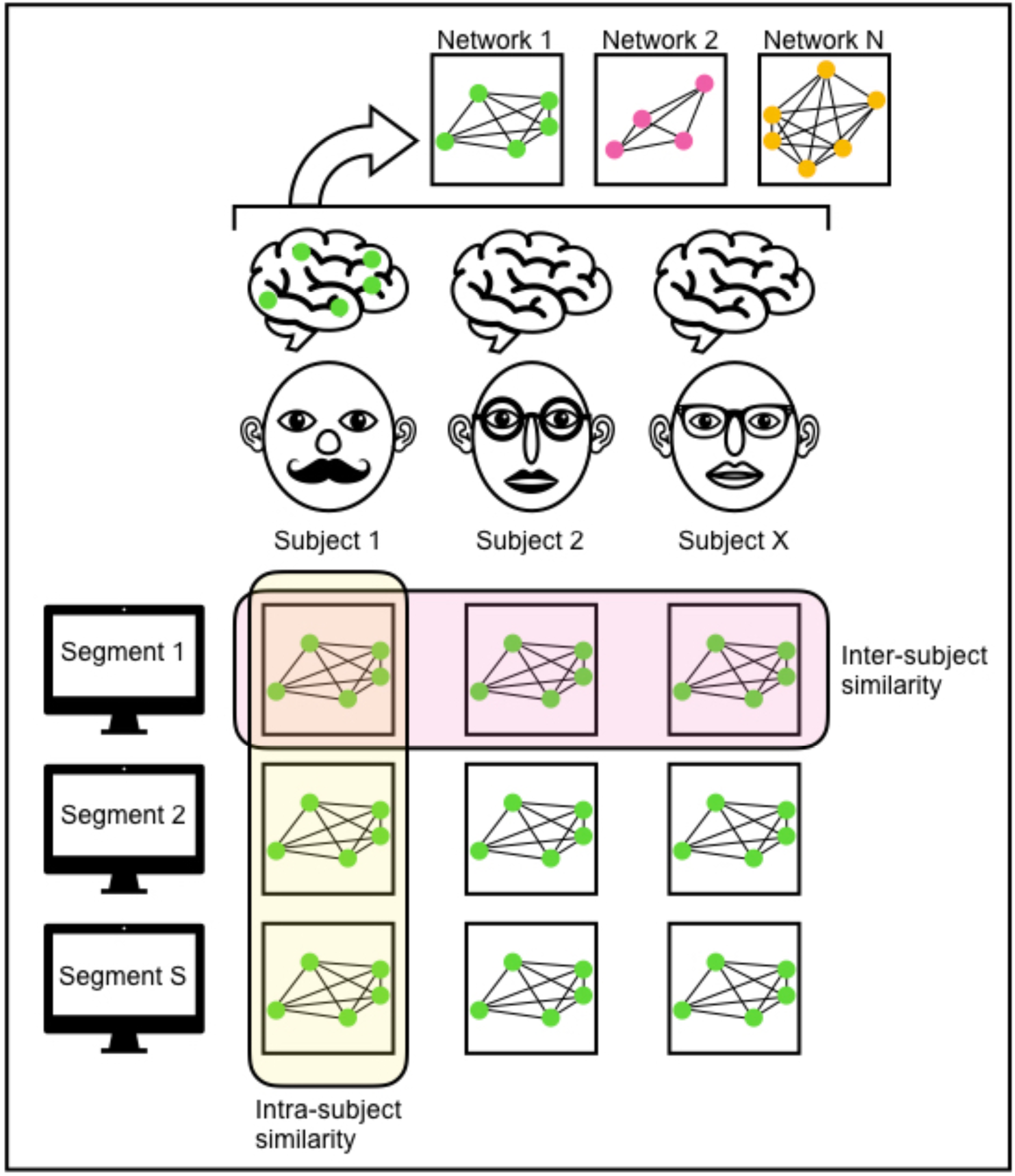
Calculation of inter– and intra-subject similarity. For each subject, functional connectomes were computed for all 14 networks in each of the 8 movie segments. Inter-subject similarity is assessed by calculating the average correlations between subjects within the same movie segments. Intra-subject similarity is assessed by calculating the average correlation between movie segments within the same subject.

### 2.5 Statistical analyses

To investigate whether portrayed emotions are different across movie segments, we conducted a one-way ANOVA for each measure. Here, IOA values were used as independent variables with the movie segments as fixed factors. Post-hoc t-tests are reported with Bonferroni-adjusted p-values.

To test whether inter-or intra-subject similarity differ across networks depending on movie segments and portrayed emotions, we applied linear mixed models (LMM) using the statsmodels python package (https://www.statsmodels.org/stable/mixed_linear.html). Specifically, we created different random intercept models by choosing network, movie segment, arousal and valence as possible fixed effects, subject identity as a random effect, and inter-or intra-subject similarity as the dependent variable. We chose subject identity as a random effect, because participants are the sampling unit of interest and contribute repeatedly to the NFC measures across all movie segments. We model individual differences by assuming different random intercepts for each subject, but no individual random slopes, because a simpler model structure was warranted by our data. Network was chosen as a fixed effect to test which networks are associated with changes in inter-or intra-subject similarity induced by NV. It was included as a categorical factor with 14 levels. Movie segment was chosen as a fixed effect to test for an effect of the length and complexity of a full-narrative movie. Portrayed valence and arousal were chosen as fixed effects to represent the emotional content of the full-narrative movie, testing if emotions portrayed in a movie affect inter-or intra-subject similarity in NFC. Models were generated using maximum likelihood to include all possible models, that is, each unique combination of one to four fixed effects and their respective interactions, resulting in 2128 models that were compared each for inter– and intra-subject similarity. The model best fitting our data was selected using Bayesian information criterion (BIC, Schwarz, 1978) and used to calculate the parameter estimates for each effect. To test whether a specific network had a significant influence on inter-or intra-subject similarity, we created a “mean network” representing the mean inter-or intra-subject similarity values across all networks that we used as a reference category to compare all other networks against. P-values were obtained using Wald tests of the best models.

## 3. Results

Emotions and an overarching narrative are hallmark features of conventional Hollywood movies, which are frequently employed in NV research because of their engaging and complex nature. However, most NV studies use only shorter clips from these movies, essentially excluding effects of the ongoing narrative. Therefore, is it not yet clear how these features might impact inter– and intra-subject similarity in NFC in a full narrative movie. Here, we investigated portrayed valence and arousal across a full narrative movie and how these factors contribute to explaining inter– and intra-subject similarity in NFC in 14 networks.

### 3.1 Movie segments and portrayed emotions

We used a previously reported description (Lab et al., 2019) of portrayed valence and arousal for comparisons between the emotional content of different movie segments. Our results showed that movie segments differed in the direction (i.e.: positive/negative valence; high/low arousal) and the extent of agreement between observers concerning these measures (Figure 2). Figure 2 shows average IOA values of each movie segment and reveals large differences in the evaluation of valence and arousal across movie segments. For segments 1, 6, 7, and 8 IOA values indicate consistency in portrayed positive valence, while the segments 4 and 5 portrayed negative valence. Segment 2 and 3 showed little consistency in the evaluation of portrayed valence, as IOA values are close to zero. Concordantly, the ANOVA on the valence IOA values resulted in a significant main effect of segment (F(7,3534) = 45.879, P < .001), and Bonferroni-corrected post-hoc testing revealed significant differences between the consecutive segments 1 and 2 (t = 3.378, p = .021), 3 and 4 (t = 7.236, p < .001), 4 and 5 (t = –3.131, p = .049), 5 and 6 (t = –8.519, p < .001) and 7 and 8 (t = 3.552, p = .011). Segment 4 had the strongest agreement on negative valence between observers, while segment 7 showed the strongest agreement on positive valence between observers. Figure 2 further shows that segments 2 and 4 portrayed high arousal, while the other segments portrayed low arousal. The ANOVA on arousal IOA values showed a significant main effect of segment as well (F(7, 3534) = 15.479, p < .001). Bonferroni-corrected post-hoc testing revealed significant differences between consecutive segments 1 and 2 (t = –13.448, p < .001), 2 and 3 (t = 10.397, p .001), 3 and 4 (t = –14.628, p < .001) and 4 and 5 (t = 10.617, p < .001).

**Figure 2.**
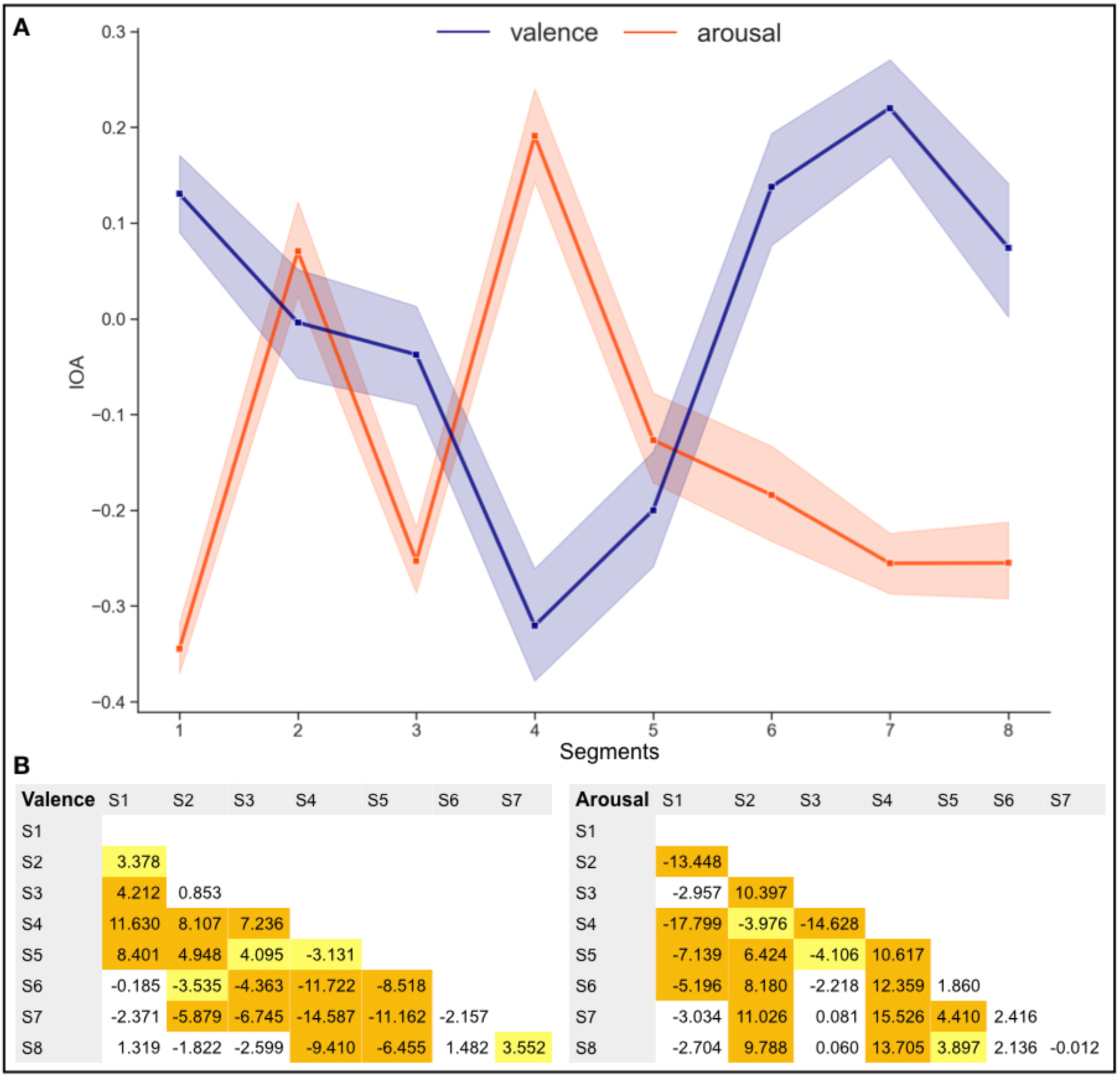
Results of the ANOVA on valence and arousal inter-observer agreement (IOA) in each movie segment. (A) Valence and arousal IOA across movie segments (portrayed valence: purple and arousal: orange). Positive IOA values indicate that observers agreed on the portrayal of positive valence and high arousal, while negative IOA values indicate that observers agreed on the portrayal of negative valence and low arousal. The amount of deviation from zero in IOA values corresponds to the strength of agreement between observers. For each movie segment, the IOA values are averaged across the entire segment. (B) Post-hoc results of the ANOVA on valence (left) and arousal (right). Bonferroni-corrected significance levels are represented by colors: orange signifies p-values < .001, yellow marks p-values < .05 and white marks no significance. S1-S8 = segments 1-8. Direction of the T-tests are column minus row element.

Given how much portrayed emotions and the narrative are intertwined, our results are best interpreted in the light of the content of the movie segments. Segment 1 spans the introduction of Forrest Gump and scenes from his childhood, containing both positive (caring mother, close friendship with neighbor girl Jenny) and negative (walking impairments, bullying) elements. Segment 2 was marked by low IOA in both valence and arousal, showing less agreement between observers on the portrayed emotions in this segment. During this segment the movie shows Forrest’s highschool and college time, addressing athletic successes and first dating experiences. Low IOA values continue in the valence dimension in segment 3, whereas observers agreed more strongly on low arousal being portrayed here. Here, the movie shows Forrest joining the army, reuniting with Jenny in a nightclub where she works as a dancer, and being deployed in the Vietnam war. Segment 4 prominently features a different pattern than any other segment: observers agreed that movie characters displayed high arousal and low valence during this segment. This can likely be attributed to the war scenes involving an ambush in Vietnam causing Forrest’s best friend’s death, and following scenes in a military hospital, although the segment also contains Forrest receiving the Medal of Honor and speaking at an anti-war rally in front of the Pentagon. Segment 5 is marked by lower IOA values indicating some negative valence and low arousal, featuring the Black Panther movement, Forrest’s ping pong career and reunions with friends Jenny and Lt. Dan. The last three segments again display a pattern of higher agreement between observers on positive valence and low arousal, when the movie spans Forrest’s successful shrimp fishing business, two episodes of living happily with Jenny, a three-year cross-country marathon, Forrest meeting his son, Jenny’s death and the ending of the movie.

### 3.2 Inter– and intra-subject similarity in NFC

Inter– and intra-subject correlations were calculated for every network on the level of single segments, i.e. the different segments of the movie. Figure 3 summarizes the results across all networks and segments based on Pearson correlation coefficients (Figure 3A), and shows results of the LMM analyses on inter-(Figure 3B) and intra-subject similarity (Figure 3C). We found that inter– and intra-subject similarity both fluctuate across time for all networks. We will further analyze the statistical significance in the following sections.

**Fig 3.**
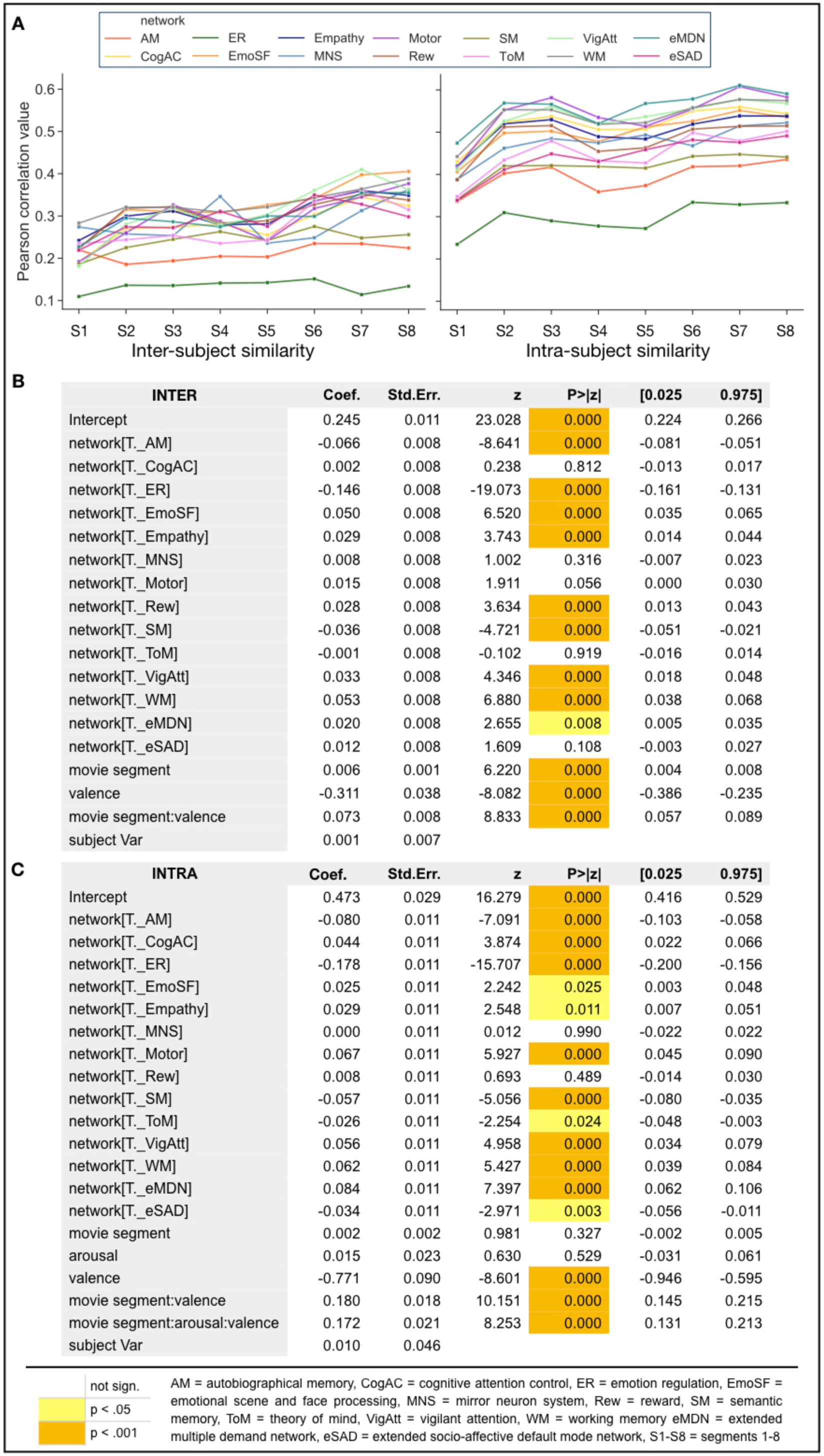
Results of the LMM showing how the fixed effect network, movie segment, arousal and valence and the random effect subject identity contribute to inter– and intra-subject similarity in NFC. (A) Inter– and intra-subject correlation of each movie segment averaged in each network, based on Pearson correlation coefficients. The x axis depicts different movie segments. The y axis represents the averaged similarity of the functional connectivity matrix derived from one subject compared to all other subjects within each network. (B) Results of the LMM on inter-subject similarity. (C) Results of the LMM on intra-subject similarity.

#### 3.2.1 Inter-subject similarity

The best model that best fitted on inter-subject similarity as selected using BIC consists of the random factor subject identity, the fixed factors network, movie segment and valence, and the interaction between the fixed factors movie segment and valence. All parameter estimates and p-values can be seen in Figure 3B. The intercept for inter-subject similarity is 0.245, representing the average inter-subject correlation value. Of all 14 networks, the AM, ER, EmoSF, Empathy, Rew, SM, VigAtt, WM and eMDN networks differed significantly from the “mean network” reference category representing the mean inter-subject similarity across all networks. The AM, ER and SM networks show negative coefficients, indicating that inter-subject similarity is lower in these networks than on average. The EmoSF, Empath, Rew, VigAtt, WM and eMDN networks were associated with higher inter-subject similarity than average. Movie segment, valence and their interaction effect reached significance as well. While movie segment and the movie segment-valence interaction were associated with higher inter-subject similarity, valence was associated with lower inter-subject similarity. The estimated coefficient for subject identity was 0.001, indicating a low effect of subject identity on inter-subject similarity and small differences between subjects.

#### 3.2.2 Intra-subject similarity

The best model that best fitted on intra-subject similarity as selected using BIC consists of the random factor subject identity, the fixed factors network, movie segment, valence and arousal, and the interactions between fixed factors movie segment and valence and between movie segment, arousal and valence. All parameter estimates and p-values can be seen in Figure 3C. The intercept for intra-subject similarity is 0.473, representing the average intra-subject correlation value. The AM, CogAC, ER, EmoSF, Empathy, Motor, SM, ToM, VigAtt, WM, eMDN and eSAD network differed significantly from the reference category representing the mean intra-subject similarity across all networks. The AM, ER, SM, ToM and eSAD networks were associated with lower intra-subject similarity, whereas the CogAC, EmoSF, Empathy, Motor, VigAtt, WM and eMDN networks were associated with higher intra-subject similarity than average. While movie segment and arousal did not reach significance, valence, the movie segment-valence interaction and the movie segment-arousal-valence interaction did. Valence was associated with lower intra-subject similarity, while the movie segment-valence and movie segment-arousal-valence interaction were associated with higher intra-subject similarity. The estimated coefficient for subject identity was 0.01, indicating a low effect of subject identity on intra-subject similarity and small differences between subjects.

## 4. Discussion

In this study, we aimed to investigate the inter– and intra-subject similarity of NFC over the course of a full narrative movie. By analyzing a publicly available dataset that contains fMRI data spanning a full narrative movie, we investigated changes in similarity over multiple, consecutive movie segments. Inter– and intra-subject similarity were best explained when accounting for network, movie segment, valence and a movie segment by valence interaction. Additionally, arousal played a role in explaining intra-subject similarity by interacting with movie segment and valence. The effect of the movie stimulus on changes in inter– and intra-subject similarity was network dependent. Comparing portrayed valence and arousal across movie segments showed that both varied across the segments, indicating differences in emotional content that we could relate to the content of the different movie segments.

### 4.1 Portrayed valence and arousal

Emotions are an important feature of movie stimuli. Shorter movies have been used to study emotion processing (Westermann et al., 1996; Carvalho et al., 2012; Schaefer at al., 2010) and longer movies might allow studying emotions across a larger timescale. Emotions are a major factor in narration (Cutting, 2016; Aldama, 2015), change over time, and dynamically interact with social context (Redcay & Moraczewski, 2019). Therefore, full narrative movies have clear advantages for studying emotions in a naturalistic setting. Additionally, emotions portrayed in movies affect inter-subject synchronization (Dziura et al., 2021) and inter-subject alignment of brain states (Chang et al., 2021), which makes them a relevant factor for studying individual differences using naturalistic viewing.

Here, we used a previously reported description (Labs et al., 2015) of portrayed valence and arousal for comparisons between the emotional content of different movie segments. Our results showed that movie segments differed in the direction (i.e.: positive/negative valence; high/low arousal) and the extent of agreement between observers concerning these measures (Figure 2). In our results a pattern emerged in which segments that were marked by high concordance in positive valence also showed good agreement in low arousal evaluations (segments 1, 6, 7, 8), whereas the reversed pattern was observable in segment 4. This indicates a potential negative relationship between valence and arousal as depicted in our movie stimuli. Valence and arousal are the bipolar dimensions in circumplex models of affect (Yik, Russell, Barrett, 1999), and the relationship between valence and arousal seems to be highly individual and related to personality and culture (Kuppens et al., 2016).

Overall, the pattern of results implies that the segments of the chosen movie stimulus differed in emotional content, which makes it valuable for inducing variability in functional networks associated with socio-emotional processing. Specifically, the Forrest Gump movie features a broad range of themes (love, friendship, politics, fate), settings (varying historical events, places, times and roles of the protagonist) and situations portraying a wide spectrum of emotions in different contexts. Our results are thus in line with studies showing that movies can elicit complex and mixed states of emotions (Schaefer et al., 2010; Carvalho et al., 2012). In particular, the Forrest Gump movie stimulus has been shown to induce distinct affective states throughout the movie, which was used to map the topographic organization of these states (Lettieri et al., 2019). Hence in accordance with the proposal by Finn and colleagues (2017), the chosen movie can evoke brain states in a meaningful manner, and thus represents a fitting stimulus for studying variability in and between subjects over time.

We investigated portrayed valence and arousal as important emotional features of the movie stimulus. Critically, Labs et al. (2015) created an annotation of the movie stimulus content, not an annotation of the viewer’s emotional experiences. In order to characterize the portrayed emotions as a relatively lower level feature, observers rated all movie scenes in randomized order to prevent “carry-over” effects from the context the scenes appear within and the current mood of the movie (Labs et al., 2015). This annotation therefore offers descriptive information about the movie stimulus rather than assessing the full emotional complexity of the movie and its effects on the viewer. The characterisation of emotion cues in single scenes offers the benefit of relating these cues to other features of the movie scenes (e.g. lighting, audio features) in future studies.

### 4.2 Inter-subject similarity

Across networks, inter-subject correlations increased over the course of the movie, indicating a general tendency of subjects’ NFC to become more similar (Figure 3A). By using LMM, we found several factors contributing to changes in inter-subject similarity, including network, movie segment, valence and interaction of movie content and valence. Specifically, the model that best explains inter-subject similarity comprises the fixed effects network, movie segment, valence and a movie segment by valence interaction, with subject identity as a random effect.

Looking more closely at the networks, we see that some networks, such as the CogAC, MNS, Motor, ToM and eSAD, do not contribute significantly to changes in inter-subject similarity. This might indicate that these networks are not sensitive to the effects of a full narrative movie and the emotions portrayed within. For those networks that are significantly modulated by movie content, we observed large variations across networks in inter-subject similarity. The AM, ER and SM networks are associated with lower inter-subject similarity, which can be seen in lower inter-subject correlation values (Figure 3A) and negative model coefficients (Figure 3B). This indicates that emotion regulation and long term memory processes are most sensitive to a full narrative movie. This might reflect the stimulus containing a highly emotional narrative and many references to real world events and history. Contrarily, the EmoSF, Empathy, Rew, VigAtt, WM and eMDN networks are associated with higher inter-subject similarity. Across all networks, the coefficients exhibit a wide range in values, with the ER network showing the highest absolute coefficient, indicating the greatest effect on inter-subject similarity. Movie segment had a small negative effect on inter-subject similarity, indicating that inter-subject similarity increases over the course of a full narrative movie, which is also reflected in a slight increase in inter-subject correlation values (Figure 3A). Previous research has shown high inter-subject variability in response to professionally produced and conventional movies that were much shorter (< 20min) than a full narrative movie (Hasson, Malach & Heeger, 2009; Vanderwal et al., 2015). It is likely that the change towards more similarity in NFC over the course of the movie results from the shared experience, which is created to evoke certain reactions and feelings in the audience. Indeed, viewers’ emotional and cognitive states can be affected and synchronized through director’s decisions, such as the camera settings, light, performance of actors, scripts and dialog, and more (Tarvainen, Westman & Oittinen, 2015; Münsterberg, 1916; Baranowski & Hecht, 2017). Studying viewers’ emotions while watching the identical stimulus used here, Lettieri et al. (2019) showed that ratings of basic emotions were consistent across viewers, indicating an overall highly similar emotional experience induced by the movie. Emphasizing the relevance of affective states in movie fMRI, previous studies showed higher alignment of brain states between subjects during highly affective events in a TV show (Chang et al., 2021) and more synchronization of amygdala activity between subjects during positive events in a “shared watching” condition (Dziura et al., 2021).

In our study, valence was associated with lower inter-subject similarity. A study by Nummenmaa et al. (2012) studied the relationship between perceived valence and arousal and inter-subject synchronization of brain activity during movie watching. They found that more negative valence was associated with increased inter-subject synchronization in an emotion-processing network and the default-mode network, while high arousal was associated with increased inter-subject synchronization in somatosensory cortices, and visual and dorsal attention networks (Nummenmaa et al., 2012). This is in line with the pattern of positive valence being associated with lower similarity in our results.

However, the movie segment by valence interaction has a positive coefficient, indicating that positive valence is associated with higher inter-subject similarity across the course of a full narrative movie. This might represent the effects of a conventional Hollywood movie orchestrating similarity in viewers’ experience by using positive portrayed emotions.

The random factor subject identity had a very small negative effect on inter-subject similarity, indicating that there were no great differences between subjects.

### 4.3 Intra-subject similarity

Our results show that intra-subject similarity increases over the course of a full narrative movie across networks (s. Figure 3A).

When selecting the best model in our LMM analysis to explain intra-subject similarity, network, movie segment, arousal, and valence emerged as relevant fixed effects. Additionally, the model included a movie segment by valence and a movie segment by arousal by valence interaction. Again, subject identity was included as a random effect.

Of all networks, only the MNS and Rew networks did not affect intra-subject similarity significantly. The AM, ER, SM, ToM and eSAD networks were associated with decreased intra-subject similarity, while the CogAC, EmoSF, Empathy, Motor, VigAtt, WM and eMDN networks were associated with increased intra-subject similarity. Similar to the results on inter-subject similarity, emotion regulation and long-term memory were most sensitive to the effects of a full narrative movie, showing the lowest intra-subject similarity across movie segments. Additionally, networks processing self– and other-related social cognition showed low intra-subject similarity, indicating that a full narrative movie might tax introspection and relating to others in a way that varies along the narrative.

Similar to the pattern of results seen in inter-subject similarity, valence was associated with lower intra-subject similarity while the movie segment by valence interaction was associated with higher intra-subject similarity. Additionally, the three-way interaction between movie segment, valence and arousal was associated with higher intra-subject similarity. This might indicate that positive valence is generally associated with lower intra-subject similarity, although the progression of movie segments and higher portrayed arousal increase intra-subject similarity. While arousal did not significantly influence inter-subject similarity, it interacts with movie segment and valence when influencing intra-subject similarity. This might indicate that arousal is a more relevant factor when investigating similarity within subjects and might prompt future comparisons on the effects of movies with different levels of arousal on single subjects. Arousal seems to be influenced by various stylistic features of a movie and can be further differentiated into subdimensions such as energetic and tense mood (Tarvainen et al., 2015).

The random factor subject identity had a very small positive effect on intra-subject similarity, indicating that there were no great differences between subjects.

### 4.5 Limitations

By using an unusually long naturalistic stimulus – a full-length movie – our study offers important insights into inter– and intra-subject similarity in NFC across a two hour acquisition period.

Our results indicate that the content of a movie is a relevant factor in naturalistic viewing, but it is not yet certain how different content or features of a movie relate to inter– and intra-subject similarity. Our study of one full narrative movie and its annotation of portrayed valence and arousal is an important first step in quantifying this relationship. To generalize our results to other movies, brain measures and samples, future research needs to expand information on available naturalistic viewing datasets (for example by creating more annotations), so that content and effects on NFC can be investigated across different datasets. It is necessary to find a good match between movies and their annotated features, methodology and research question (Eickhoff et al., 2020; Saarimäki, 2021; Grall & Finn, 2022).

Our study comes at the cost of investigating only the effects of a single movie. Comparisons with an equally long resting state acquisition or a movie stimulus without a narrative would have given stronger evidence for the effect of full narrative movies. However, there were no such scans available in this dataset. Analyses of additional full narrative movies might expand the insights gained into the effects of different narratives. The choice of a conventional Hollywood movie might have led to higher inter-subject similarity (Hasson, Malach, Heeger, 2009; Vanderwal et al., 2015; Chang et al., 2021; Tarvainen, Westman & Oittinen, 201; Baranowski & Hecht, 2017), while more ambiguous or emotionally and socially equivocal movies could enhance inter-individual differences to a greater degree. Familiarity with a movie stimulus has been discussed as a potential factor for driving individual differences. An effect of repeated movie watching in functional connectivity on the network level has been shown before (Andric et al., 2016). However, such effects can be assumed to be low in our sample. All participants were familiar with the narrative of the movie, and only one participant reported to never have seen the movie (Hanke et al., 2014).

Given the unusual length of data acquisition, effects of the MRI measurement might have influenced the participants’ focus on and perception of the movie stimulus. For example, participants might have needed some time to familiarize themselves with the MRI scanner. However, as all participants had already participated in previous MRI measurements of the studyforrest project (Hanke et al., 2016; Sengupta et al., 2016), high familiarity to MRI scanning and all related procedures was present in this sample. The length of acquisition might also have affected the participants’ attention. Previous studies indicate that movie watching is very engaging and might decrease drowsiness and sleep in the scanner (Eickhoff, Milham & Vanderwal, 2020), but attention might still have been impacted over such a long duration. NV paradigms are designed to include minimal participant instructions so as not to influence participants’ perception of the stimuli or add task demands not directly related to movie watching. In future studies, post-hoc questionnaires might be useful to estimate attention fluctuations, distractions, drowsiness and other potential confounds that might have occurred during data acquisition.

The analyses of this study focussed on the approximately 15-minute segments the data was acquired in, splitting the movie into 8 segments. Time windows for analysis of NV data have varied in the literature and optimal time windows and scan lengths are still debatable. Uri Hasson’s work on temporal receptive windows focuses on window sizes on the level of seconds (e.g. time windows of ∼4 (“short”), ∼12 (“intermediate”), and ∼36 seconds (“long”)) (Hasson et al., 2010). Based on naturalistic viewing data, single subject identification accuracy was positively impacted by longer scan durations (Vanderwal et al., 2017; scan duration with highest accuracy ranged from ∼4.5 to ∼7 min depending on movie stimulus) and movies of ∼2.5 min length can be sufficient for behavioral prediction (Finn & Bandettini, 2021). Efforts for providing normative data during movie watching have been recommended to use minimally 10 min and optimally at least 25 min duration per movie (Eickhoff et al., 2020). Irrespective of naturalistic viewing, reliability of functional connectivity measures increases with time, with indications that less than 10 min of RS scan duration may not capture functional connectivity features reliably (Laumann et al., 2015, 9 to 27 min durations; Noble et al., 2017, 5 to 25 min durations). These examples show that optimal scan duration may depend strongly on the research question at hand, with advantages coming from longer durations. In our study, employing a FNM with the focus on a continuously unfolding and dynamic narrative might speak for longer scan durations to capture the effects of these “longer term” story dynamics. While the long fMRI acquisition spanning a FNM is a great advantage to our study, it comes with the disadvantage of a small sample size. Replication in other datasets is an important next step, although this specific dataset remains unique in its stimulus and annotations.

The number of nodes constituting each meta-analytical network was different between networks used in this study. Recently, the influence of network size on single subject identifiability based on NV data has been investigated (Kröll et al., 2023), indicating that the number of nodes in a network are a relevant factor in network-level analyses. The networks that were used in this study are based on meta-analysis and represent various cognitive and psychological domains, so that network size is inherent to each network and cannot be adapted at will.

This study used the preprocessed data made available by the original authors of the dataset (Hanke et al., 2016). We acknowledge that further preprocessing steps, such as scrubbing, might influence the results. However, data quality control of the original dataset authors revealed very few motion artifacts, highlighting the beneficial effect of movie watching on participant motion (Hanke et al., 2016).

### 4.6 Conclusion

The present study is the first to investigate inter– and intra-subject similarity in NFC across a full narrative movie. Our results show that inter– and intra-subject similarity in NFC were sensitive to the progressing narrative and emotions portrayed in the movie. The emotion regulation network displayed the lowest similarity within and between subjects in NFC, followed by networks associated with long-term memory processing. The sensitivity of these networks to the full narrative movie might be explained by the highly emotional narrative and continuous references to real world historical events, highlighting the importance of specific features and content of the chosen movie stimulus. The overarching narrative gives a unique possibility to study emotions in a social context and how they develop over time. These socio-cognitive aspects seem to specifically influence similarity within subjects, as low intra-subject similarity was additionally seen in networks involved in self– and other-related cognition. Altogether, these results show that a network perspective might help to elucidate the effects of different movie stimuli on specific cognitive domains. Characterizing movie stimuli in more detail to explore the effects of different features on inter– and intra-subject similarity is critical for future research in naturalistic viewing.

## Acknowledgements

The neuroimaging data was downloaded from and is openly available at https://github.com/psychoinformatics-de/studyforrest-data-phase2. The code used by the original authors of the dataset to minimally preprocess the neuromaging data can be found under https://github.com/psychoinformatics-de/studyforrest-data-aligned/tree/master/code. The valence and arousal inter observer agreement measures were downloaded from and are available at https://f1000research.com/articles/4-92/v1#DS0. The code to divide inter observer agreement time series according to the movie segments was downloaded from and is available at https://osf.io/tzpdf/.

The work was supported by: Deutsche Forschungsgemeinschaft (DFG), National Institute of Mental Health (R01-MH074457), Helmholtz Portfolio Theme “Supercomputing and Modeling for the Human Brain”, European Union’s Horizon 2020 Research, Innovation, Programme under Grant Agreement No. 945539 (HBP SGA3). Open access publication funded by the DFG – 491111487.

The authors declare no conflict of interest.

The Ethics committee of Otto-Von-Guericke University, Germany approved acquisition of the data in the “studyforrest” project.

## References

1. Adolphs, R., Nummenmaa, L., Todorov, A., & Haxby, J. V. (2016). Data-driven approaches in the investigation of social perception. Philosophical Transactions of the Royal Society B: Biological Sciences, 371(1693). 10.1098/rstb.2015.0367

2. Amft, M., Bzdok, D., Laird, A. R., Fox, P. T., Schilbach, L., & Eickhoff, S. B. (2015). Definition and characterization of an extended social-affective default network. Brain Structure and Function, 220(2), 1031–1049. 10.1007/s00429-013-0698-0

3. Andric, M., Goldin-Meadow, S., Small, S. L., & Hasson, U. (2016). Repeated movie viewings produce similar local activity patterns but different network configurations. NeuroImage, 142, 613–627. 10.1016/j.neuroimage.2016.07.061

4. Baranowski, A., & Hecht, H. (2017). One Hundred Years of Photoplay: Hugo Münsterberg’s Lasting Contribution to Cognitive Movie Psychology. Projections, 11(2), 1–21. 10.3167/proj.2017.110202

5. Binder, J. R., & Desai, R. H. (2011). The Neurobiology of Semantic Memory Jeffrey. Trends in Cognitive Sciences, 15(11), 527–536. 10.1016/j.tics.2011.10.001.The

6. Buhle, J. T., Silvers, J. A., Wage, T. D., Lopez, R., Onyemekwu, C., Kober, H., … Ochsner, K. N. (2014). Cognitive reappraisal of emotion: A meta-analysis of human neuroimaging studies. Cerebral Cortex, 24(11), 2981–2990. 10.1093/cercor/bht154

7. Bzdok, D., Schilbach, L., Vogeley, K., Schneider, K., Laird, A. R., Langner, R., & Eickhoff, S. B. (2012). Parsing the neural correlates of moral cognition: ALE meta-analysis on morality, theory of mind, and empathy. Brain Structure and Function, 217(4), 783–796. 10.1007/s00429-012-0380-y

8. Camilleri, J. A., Müller, V. I., Fox, P., Laird, A. R., Hoffstaedter, F., Kalenscher, T., & Eickhoff, S. B. (2018). Definition and characterization of an extended multiple-demand network. NeuroImage, 165(October 2017), 138–147. 10.1016/j.neuroimage.2017.10.020

9. Carvalho, S., Leite, J., Galdo-Álvarez, S., & Gonçalves, Ó. F. (2012). The emotional movie database (EMDB): A self-report and psychophysiological study. Applied Psychophysiology Biofeedback, 37(4), 279–294. 10.1007/s10484-012-9201-6

10. Caspers, S., Zilles, K., Laird, A. R., & Eickhoff, S. B. (2010). ALE meta-analysis of action observation and imitation in the human brain. NeuroImage, 50(3), 1148–1167. 10.1016/j.neuroimage.2009.12.112

11. Chang, L. J., Jolly, E., Cheong, J. H., Rapuano, K. M., Greenstein, N., Chen, P. H. A., & Manning, J. R. (2021). Endogenous variation in ventromedial prefrontal cortex state dynamics during naturalistic viewing reflects affective experience. Science Advances, 7(17). 10.1126/sciadv.abf7129

12. Cieslik, E. C., Mueller, V. I., Eickhoff, C. R., Langner, R., & Eickhoff, S. B. (2015). Three key regions for supervisory attentional control: Evidence from neuroimaging meta-analyses. Neuroscience and Biobehavioral Reviews, 48, 22–34. 10.1016/j.neubiorev.2014.11.003

13. Delgado-Herrera, M., Reyes-Aguilar, A., & Giordano, M. (2021). What Deception Tasks Used in the Lab Really Do: Systematic Review and Meta-analysis of Ecological Validity of fMRI Deception Tasks. Neuroscience, 468, 88–109. 10.1016/j.neuroscience.2021.06.005

14. Demirtaş, M., Ponce-Alvarez, A., Gilson, M., Hagmann, P., Mantini, D., Betti, V., … Deco, G. (2019). Distinct modes of functional connectivity induced by movie-watching. NeuroImage, 184(June 2018), 335–348. 10.1016/j.neuroimage.2018.09.042

15. Dosenbach, N. U. F., Visscher, K. M., Palmer, E. D., Miezin, F. M., Wenger, K. K., Kang, H. C., … Petersen, S. E. (2006). A Core System for the Implementation of Task Sets. Neuron, 50(5), 799–812. 10.1016/j.neuron.2006.04.031

16. Dziura, S. L., Merchant, J. S., Alkire, D., Rashid, A., Shariq, D., Moraczewski, D., & Redcay, E. (2021). Effects of social and emotional context on neural activation and synchrony during movie viewing. Human Brain Mapping, 42(18), 6053–6069. 10.1002/hbm.25669

17. Eickhoff, S. B., Bzdok, D., Laird, A. R., Kurth, F., & Fox, P. T. (2012). Activation likelihood estimation revisited. NeuroImage, 59(3), 2349–2361. 10.1016/j.neuroimage.2011.09.017.Activation

18. Eickhoff, Simon B., & Grefkes, C. (2011). Approaches for the integrated analysis of structure, function and connectivity of the human brain. Clinical EEG and Neuroscience, 42(2), 107–121. 10.1177/155005941104200211

19. Eickhoff, Simon B., Laird, A. R., Grefkes, C., Wang, L. E., Zilles, K., & Fox, P. T. (2009). Coordinate-based activation likelihood estimation meta-analysis of neuroimaging data: A random-effects approach based on empirical estimates of spatial uncertainty. Human Brain Mapping, 30(9), 2907–2926. 10.1002/hbm.20718

20. Eickhoff, Simon B., Milham, M., & Vanderwal, T. (2020). Towards clinical applications of movie fMRI. NeuroImage, 217. 10.1016/j.neuroimage.2020.116860

21. Elliott, M. L., Knodt, A. R., Cooke, M., Kim, M. J., Melzer, T. R., Keenan, R., … Hariri, A. R. (2019). General functional connectivity: Shared features of resting-state and task fMRI drive reliable and heritable individual differences in functional brain networks. NeuroImage, 189(January), 516–532. 10.1016/j.neuroimage.2019.01.068

22. Etkin, A., Büchel, C., & Gross, J. J. (2015). The neural bases of emotion regulation. Nature Reviews Neuroscience, 16(11), 693–700. 10.1038/nrn4044

23. Finn, E. S., & Bandettini, P. A. (2021). Movie-watching outperforms rest for functional connectivity-based prediction of behavior. NeuroImage, 235(March), 117963. 10.1016/j.neuroimage.2021.117963

24. Finn, E. S., Scheinost, D., Finn, D. M., Shen, X., Papademetris, X., & Constable, R. T. (2017). Can brain state be manipulated to emphasize individual differences in functional connectivity? NeuroImage, 160(March), 140–151. 10.1016/j.neuroimage.2017.03.064

25. Fox, K. C. R., Spreng, R. N., Ellamil, M., Andrews-Hanna, J. R., & Christoff, K. (2015). The wandering brain: Meta-analysis of functional neuroimaging studies of mind-wandering and related spontaneous thought processes. NeuroImage, 111, 611–621. 10.1016/j.neuroimage.2015.02.039

26. Fox, M. D., Snyder, A. Z., Vincent, J. L., Corbetta, M., Van Essen, D. C., & Raichle, M. E. (2005). The human brain is intrinsically organized into dynamic, anticorrelated functional networks. Proceedings of the National Academy of Sciences of the United States of America, 102(27), 9673–9678. 10.1073/pnas.0504136102

27. Gonzalez-Castillo, J., Kam, J. W. Y., Hoy, C. W., & Bandettini, P. A. (2021). How to interpret resting-state fMRI: ask your participants. Journal of Neuroscience, 41(6), 1130–1141.

28. Grall, C., & Finn, E. S. (2022). Leveraging the power of media to drive cognition: A media-informed approach to naturalistic neuroscience. Social Cognitive and Affective Neuroscience, 17(6), 598–608. 10.1093/scan/nsac019

29. Gross, J. J. (2015). Emotion Regulation: Current Status and Future Prospects. Psychological Inquiry, 26(1), 1–26. 10.1080/1047840X.2014.940781

30. Gross, J. J., & Levenson, R. W. (1995). Emotion Elicitation using Films. Cognition and Emotion, 9(1), 87–108. 10.1080/02699939508408966

31. Hanke, M., Adelhöfer, N., Kottke, D., Iacovella, V., Sengupta, A., Kaule, F. R., … Stadler, J. (2016). A studyforrest extension, simultaneous fMRI and eye gaze recordings during prolonged natural stimulation. Scientific Data, 3, 160092. 10.1038/sdata.2016.92

32. Hanke, Michael, Baumgartner, F. J., Ibe, P., Kaule, F. R., Pollmann, S., Speck, O., … Stadler, J. (2014). A high-resolution 7-Tesla fMRI dataset from complex natural stimulation with an audio movie. Scientific Data, 1, 1–18. 10.1038/sdata.2014.3

33. Hasson, U., Landesman, O., Knappmeyer, B., Vallines, I., Rubin, N., & Heeger, D. J. (2008). Neurocinematics: The Neuroscience of Film. Projections, 2(1), 1–26. 10.3167/proj.2008.020102

34. Hasson, U., Malach, R., & Heeger, D. J. (2010). Reliability of cortical activity during natural stimulation. Trends in Cognitive Sciences, 14(1), 40–48. 10.1016/j.tics.2009.10.011

35. Hasson, U., Nir, Y., Levy, I., Fuhrmann, G., & Malach, R. (2004). Intersubject Synchronization of Cortical Activity During Natural Vision. Science, 303(MARCH), 1634–1640. 10.1126/science.1089506

36. Igelström, K. M., & Graziano, M. S. A. (2017). The inferior parietal lobule and temporoparietal junction: A network perspective. Neuropsychologia, 105(December 2016), 70–83. 10.1016/j.neuropsychologia.2017.01.001

37. Kim, D., Cho, Y., & Park, K. S. (2018). Comparative analysis of affective and physiological responses to emotional movies. Human-Centric Computing and Information Sciences, 8(1). 10.1186/s13673-018-0138-5

38. Kingstone, A., Smilek, D., & Eastwood, J. D. (2008). Cognitive Ethology: A new approach for studying human cognition. British Journal of Psychology, 99(3), 317–340. 10.1348/000712607X251243

39. Kröll, J. P., Friedrich, P., Li, X., Patil, K. R., Mochalski, L., Waite, L., … Weis, S. (2023). Naturalistic viewing increases individual identifiability based on connectivity within functional brain networks. NeuroImage, 273(April), 120083. 10.1016/j.neuroimage.2023.120083

40. Kuppens, P., Tuerlinckx, F., Yik, M., Koval, P., Coosemans, J., Zeng, K. J., & Russell, J. A. (2017). The Relation Between Valence and Arousal in Subjective Experience Varies With Personality and Culture. Journal of Personality, 85(4), 530–542. 10.1111/jopy.12258

41. Labs, A., Reich, T., Schulenburg, H., Boennen, M., Mareike, G., Golz, M., … Hanke, M. (2015). Portrayed emotions in the movie “Forrest Gump” [version 1; referees: 2 approved] Referee Status:, (0). 10.12688/f1000research.6230.1

42. Langner, R., & Eickhoff, S. B. (2013). Sustaining attention to simple tasks: A meta-analytic review of the neural mechanisms of vigilant attention. Psychological Bulletin, 139(4), 870–900. 10.1037/a0030694

43. Laumann TO, Snyder AZ, Mitra A, Gordon EM, Gratton C, Adeyemo B, Gilmore AW, Nelson SM, Berg JJ, Greene DJ, McCarthy JE, Tagliazucchi E, Laufs H, Schlaggar BL, Dosenbach NUF, Petersen SE. On the Stability of BOLD fMRI Correlations. Cereb Cortex. 2017 Oct 1;27(10):4719–4732. doi: 10.1093/cercor/bhw265. PMID: 27591147; PMCID: PMC6248456.

44. Lerner, Y., Honey, C. J., Silbert, L. J., & Hasson, U. (2011). Topographic mapping of a hierarchy of temporal receptive windows using a narrated story. Journal of Neuroscience, 31(8), 2906–2915. 10.1523/JNEUROSCI.3684-10.2011

45. Lettieri, G., Handjaras, G., Ricciardi, E., Leo, A., Papale, P., Betta, M., … Cecchetti, L. (2019). Emotionotopy in the human right temporo-parietal cortex. Nature Communications, 10(1). 10.1038/s41467-019-13599-z

46. Liu, X., Hairston, J., Schrier, M., & Fan, J. (2011). Common and distinct networks underlying reward valence and processing stages, 35(5), 1219–1236. 10.1016/j.neubiorev.2010.12.012.Common

47. Margulies, D. S., Ghosh, S. S., Goulas, A., Falkiewicz, M., Huntenburg, J. M., Langs, G., … Smallwood, J. (2016). Situating the default-mode network along a principal gradient of macroscale cortical organization. Proceedings of the National Academy of Sciences of the United States of America, 113(44), 12574–12579. 10.1073/pnas.1608282113

48. Meconi, F., Linde-Domingo, J. S. Ferreira, C., Michelmann, S., Staresina, B., Apperly, I. A., & Hanslmayr, S. (2021). EEG and fMRI evidence for autobiographical memory reactivation in empathy. Human Brain Mapping, 42(14), 4448–4464. 10.1002/hbm.25557

49. Messel, M. S., Raud, L., Hoff, P. K., Skaftnes, C. S., & Huster, R. J. (2019). Strategy switches in proactive inhibitory control and their association with task-general and stopping-specific networks. Neuropsychologia, 135, 107220. 10.1016/j.neuropsychologia.2019.107220

50. Morawetz, C., Bode, S., Baudewig, J., & Heekeren, H. R. (2017). Effective amygdala-prefrontal connectivity predicts individual differences in successful emotion regulation. Social Cognitive and Affective Neuroscience, 12(4), 569–585. 10.1093/scan/nsw169

51. Murphy, K., Bodurka, J., & Bandettini, P. A. (2007). How long to scan? The relationship between fMRI temporal signal to noise ratio and necessary scan duration. NeuroImage, 34(2), 565–574. 10.1016/j.neuroimage.2006.09.032

52. Nastase, S. A., Gazzola, V., Hasson, U., & Keysers, C. (2019). Measuring shared responses across subjects using intersubject correlation. Social Cognitive and Affective Neuroscience, 14(6), 669–687. 10.1093/scan/nsz037

53. Noble S, Scheinost D, Finn ES, Shen X, Papademetris X, McEwen SC, Bearden CE, Addington J, Goodyear B, Cadenhead KS, Mirzakhanian H, Cornblatt BA, Olvet DM, Mathalon DH, McGlashan TH, Perkins DO, Belger A, Seidman LJ, Thermenos H, Tsuang MT, van Erp TGM, Walker EF, Hamann S, Woods SW, Cannon TD, Constable RT. Multisite reliability of MR-based functional connectivity. Neuroimage. 2017 Feb 1;146:959–970. doi: 10.1016/j.neuroimage.2016.10.020. Epub 2016 Oct 13. PMID: 27746386; PMCID: PMC5322153.

54. Nostro, A. D., Müller, V. I., Varikuti, D. P., Pläschke, R. N., Hoffstaedter, F., Langner, R., … Eickhoff, S. B. (2018). Predicting personality from network-based resting-state functional connectivity. Brain Structure and Function, 223(6), 2699–2719. 10.1007/s00429-018-1651-z

55. Nummenmaa, L., Glerean, E., Viinikainen, M., Jääskeläinen, I. P., Hari, R., & Sams, M. (2012). Emotions promote social interaction by synchronizing brain activity across individuals. Proceedings of the National Academy of Sciences of the United States of America, 109(24), 9599–9604. 10.1073/pnas.1206095109

56. Pervaiz, U., Vidaurre, D., Woolrich, M. W., & Smith, S. M. (2020). Optimising network modelling methods for fMRI. NeuroImage, 211(November 2019), 116604. 10.1016/j.neuroimage.2020.116604

57. Pläschke, R. N., Cieslik, E. C., Müller, V. I., Hoffstaedter, F., Plachti, A., Varikuti, D. P., … Eickhoff, S. B. (2017). On the integrity of functional brain networks in schizophrenia, Parkinson’s disease, and advanced age: Evidence from connectivity-based single-subject classification. Human Brain Mapping, 38(12), 5845–5858. 10.1002/hbm.23763

58. Power, J. D., Cohen, A. L., Nelson, S. M., Wig, G. S., Barnes, K. A., Church, J. A., … Petersen, S. E. (n.d.). Functional network organization of the human brain. Bone, 72(4), 665–678. 10.1016/j.neuron.2011.09.006

59. Reggente, N., Essoe, J. K. Y., Aghajan, Z. M., Tavakoli, A. V., McGuire, J. F., Suthana, N. A., & Rissman, J. (2018). Enhancing the ecological validity of fMRI memory research using virtual reality. Frontiers in Neuroscience, 12(JUN), 1–9. 10.3389/fnins.2018.00408

60. Rottschy, C., Langner, R., Dogan, I., Reetz, K., Laird, A. R., Schulz, J. B., … Eickhoff, S. B. (2012). Modelling neural correlates of working memory: A coordinate-based meta-analysis. NeuroImage, 60(1), 830–846. 10.1016/j.neuroimage.2011.11.050

61. Sabatinelli, D., Fortune, E. E., Li, Q., Siddiqui, A., Krafft, C., Oliver, W. T., … Jeffries, J. (2011). Emotional perception: Meta-analyses of face and natural scene processing. NeuroImage, 54(3), 2524–2533. 10.1016/j.neuroimage.2010.10.011

62. Saarimäki, H. (2021). Naturalistic Stimuli in Affective Neuroimaging: A Review. Frontiers in Human Neuroscience, 15(June). 10.3389/fnhum.2021.675068

63. Schaefer, Alexander, Kong, R., Gordon, E. M., Laumann, T. O., Zuo, X.-N., Holmes, A. J., … Yeo, B. T. T. (2017). Local-Global Parcellation of the Human Cerebral Cortex from Intrinsic Functional Connectivity MRI. *Cerebral Cortex*, (July 2017), 1–20. 10.1093/cercor/bhx179

64. Schaefer, Alexandre, Nils, F., Philippot, P., & Sanchez, X. (2010). Assessing the effectiveness of a large database of emotion-eliciting films: A new tool for emotion researchers. Cognition and Emotion, 24(7), 1153–1172. 10.1080/02699930903274322

65. Schwarz, G. (1982). Estimating the Dimension of a Model. Annals of Statistics, 6(2), 461–464.

66. Sengupta, A., Kaule, F. R., Guntupalli, J. S., Hoffmann, M. B., Häusler, C., Stadler, J., & Hanke, M. (2016). A studyforrest extension, retinotopic mapping and localization of higher visual areas. Scientific Data, 3, 1–14. 10.1038/sdata.2016.93

67. Simony, E., Honey, C. J., Chen, J., Lositsky, O., Yeshurun, Y., Wiesel, A., & Hasson, U. (2016). Dynamic reconfiguration of the default mode network during narrative comprehension. Nature Communications, 7(May 2015), 1–13. 10.1038/ncomms12141

68. Skerry, A. E., & Saxe, R. (2014). A common neural code for perceived and inferred emotion. Journal of Neuroscience, 34(48), 15997–16008. 10.1523/JNEUROSCI.1676-14.2014

69. Smith, S. M., Fox, P. T., Miller, K. L., Glahn, D. C., Fox, P. M., Mackay, C. E., … Beckmann, C. F. (2009). Correspondence of the brain’s functional architecture during activation and rest. Proceedings of the National Academy of Sciences of the United States of America, 106(31), 13040–13045. 10.1073/pnas.0905267106

70. Spiers, H. J., & Maguire, E. A. (2007). Decoding human brain activity during real-world experiences. Trends in Cognitive Sciences, 11(8), 356–365. 10.1016/j.tics.2007.06.002

71. Spreng, R. N., & Grady, C. L. (2010). Patterns of brain activity supporting autobiographical memory, prospection, and theory of mind, and their relationship to the default mode network. Journal of Cognitive Neuroscience, 22(6), 1112–1123. 10.1162/jocn.2009.21282

72. Tarvainen, J., Westman, S., & Oittinen, P. (2015). The way films feel: Aesthetic features and mood in film. *Psychology of Aesthetics*, Creativity, and the Arts, 9(3), 254–265. 10.1037/a0039432

73. Thomas Yeo, B. T., Krienen, F. M., Sepulcre, J., Sabuncu, M. R., Lashkari, D., Hollinshead, M., … Buckner, R. L. (2011). The organization of the human cerebral cortex estimated by intrinsic functional connectivity. Journal of Neurophysiology, 106(3), 1125–1165. 10.1152/jn.00338.2011

74. Van Atteveldt, N., Van Kesteren, M. T. R., Braams, B., & Krabbendam, L. (2018). Neuroimaging of learning and development: Improving ecological validity. Frontline Learning Research, 6(3), 186–203. 10.14786/flr.v6i3.366

75. Vanderwal, T., Eilbott, J., & Castellanos, F. X. (2019). Movies in the magnet: Naturalistic paradigms in developmental functional neuroimaging. Developmental Cognitive Neuroscience, 36(October 2018), 100600. 10.1016/j.dcn.2018.10.004

76. Vanderwal, T., Eilbott, J., Finn, E. S., Craddock, R. C., Turnbull, A., & Castellanos, F. X. (2017). Individual differences in functional connectivity during naturalistic viewing conditions. NeuroImage, 157(June), 521–530. 10.1016/j.neuroimage.2017.06.027

77. Vanderwal, T., Kelly, C., Eilbott, J., Mayes, L. C., & Castellanos, F. X. (2015). Inscapes: A movie paradigm to improve compliance in functional magnetic resonance imaging. NeuroImage, 122, 222–232. 10.1016/j.neuroimage.2015.07.069

78. Visconti Di Oleggio Castello, M., Halchenko, Y. O., Guntupalli, J. S., Gors, J. D., & Gobbini, M. I. (2017). The neural representation of personally familiar and unfamiliar faces in the distributed system for face perception. Scientific Reports, 7(1), 1–14. 10.1038/s41598-017-12559-1

79. Westermann, R., Spies, K., Stahl, G., & Hesse, F. W. (1996). Relative effectiveness and validity of mood induction procedures: a meta-analysis RAINER. European Journal of Social Psychology, 26, 557–580.

80. Witt, S. T., Meyerand, M. E., & Laird, A. R. (2008). Functional neuroimaging correlates of finger tapping task variations: An ALE meta-analysis, 71(2), 233–236. 10.1038/mp.2011.182.doi

81. Yik, M. S. M., Russell, J. A., & Barrett, L. F. (1999). Structure of self-reported current affect: Integration and beyond. Journal of Personality and Social Psychology, 77(3), 600–619. 10.1037/0022-3514.77.3.600

