## Supplementary Material for "Inter- and intra-subject similarity in network functional connectivity across a full narrative movie"

### Supplement

| participant_id | gender | age |
|---|---|---|
| sub-01 | m | 30-35 |
| sub-02 | m | 30-35 |
| sub-03 | f | 20-25 |
| sub-04 | f | 20-25 |
| sub-05 | m | 25-30 |
| sub-06 | m | 20-25 |
| sub-09 | m | 30-35 |
| sub-10 | f | 20-25 |
| sub-14 | f | 30-35 |
| sub-15 | m | 25-30 |
| sub-16 | m | 35-40 |
| sub-17 | m | 30-35 |
| sub-18 | m | 30-35 |
| sub-19 | f | 20-25 |
| sub-20 | f | 25-30 |

Table S1. Information on participant ID, gender and age range as provided in https://openneuro.org/datasets/ds000113/versions/1.3.0.

S2 Functional Brain Networks

Autobiographical memory (AM)

A central domain involved in a core network linked to self-projection and scene construction is remembering personal events from one's own past (Spreng, Mar, & Kim, 2008). Within that core network, an autobiographical memory network was found within the medial and lateral temporal cortices, precuneus, posterior cingulate cortex, retrosplenial cortex, temporo-parietal junction, lateral prefrontal and occipital cortices and medial prefrontal cortex (Spreng et al., 2008).

Cognitive attention control (CogAC)

The CogAC network (Cieslik, Mueller, Eickhoff, Langner, & Eickhoff, 2013) is involved in higher level control processes and goal-oriented behaviour and consists of the anterior insula, inferior frontal gyrus, dorsolateral prefrontal cortex, dorsal premotor cortex, bilateral intraparietal sulcus and superior parietal lobe, right temporo-parietal junction, left inferior occipital gyrus, pre-supplementary motor area and anterior midcingulate cortex, as well as the right thalamus and right caudate nucleus.

Extended multiple demand network (eMDN)

Executive functions are fundamental to a variety of behaviours and recruit a core network of brain regions linked to a multiple demand system and additional more task-specific sub-networks. This eMDN consists of the bilateral inferior frontal gyrus, insula, supplementary motor area, intraparietal sulcus, middle frontal gyrus, dorsal pre-motor cortex, putamen, thalamus and the left inferior temporal gyrus (Camilleri et al., 2016).

Emotional scene and face processing (EmoSF)

FMRI studies on emotional processing often make use of visual, emotional face or scene stimuli (Sabatinelli et al., 2011). A meta-analysis on brain regions involved in the visual perception of emotional faces or scenes revealed consistent activation in the medial prefrontal cortex, bilateral inferior, middle and superior frontal gyrus, amygdala, parahippocampal gyrus, fusiform gyrus, medial prefrontal gyrus, orbitofrontal gyrus, lateral occipital cortex, thalamus, pulvinar, right middle temporal gyrus and right anterior cingulate cortex (Sabatinelli et al., 2011).

Empathy

Empathy is the adoption of another's emotional state (Singer & Lamm, 2009). Investigating its involvement in moral decision-making, a meta-analysis revealed an empathy network consisting of the bilateral dorsomedial prefrontal cortex, anterior insula, inferior frontal gyrus, supplementary motor area, cingulate cortex, temporo-parietal junction, right amygdala, right middle temporal gyrus, right posterior superior temporal sulcus, left anterior thalamus, right posterior thalamus, right hippocampus, midbrain and right pallidum (Bzdok et al., 2012).

Theory of mind (ToM)

The same meta-analysis additionally investigated theory of mind, the ability to contemplate another’s thoughts, desires and behaviour (Premack and Woodruff, 1978; Frith and Frith, 2003). Brain regions involved in theory of mind were the ventromedial and dorsomedial prefrontal cortex, frontopolar cortex, precuneus, bilateral temporo-parietal junction, temporal pole, middle temporal gyrus, posterior superior temporal sulcus, inferior frontal gyrus and right visual area V5 (Bzdok et al., 2012).

Emotion regulation (ER)

Cognitive reappraisal, which refers to changing one’s interpretation of affective stimuli, is an important strategy of emotion regulation (Buhle et al., 2014). A meta-analysis identified the following brain regions as consistently involved in the support or moderation of cognitive reappraisal: the bilateral inferior frontal gyrus, middle frontal gyrus, superior parietal lobe, amygdala, right medial frontal gyrus, left anterior cingulate gyrus, left anterior insula, left superior temporal gyrus and left middle temporal gyrus (Buhle et al., 2014).

Extended socio-affective default network (eSAD)

The default mode network (Gusnard and Raichle, 2001 -> check REF) is closely linked to socio-affective processing (Schilbach et al., 2012 -> check REF). Therefore, Amft et al. (2015) performed a conjunction analysis on regions involved both in the DMN and social or affective tasks, resulting in the eSAD. This network consists of the anterior cingulate cortex, bilateral amygdala and hippocampus, temporo-parietal junction, ventral basal ganglia, precuneus, subgenual cingulate cortex, ventromedial and dorsomedial prefrontal cortex and left middle temporal gyrus and sulcus (Amft et al., 2015).

Mirror neuron system (MNS)

Action observation and imitation tasks can give insight into the human mirror neuron system while avoiding invasive single-cell recordings used to study mirror neurons in non-human primates (Caspers, Zilles, Laird, & Eickhoff, 2016). A meta-analysis on action observation and imitation revealed an underlying network consisting of the bilateral inferior frontal gyrus, primary somatosensory cortex, lateral occipital lobe, right fusiform face and body area, left medial premotor cortex, left posterior middle temporal gyrus and right superior parietal lobe (Caspers et al., 2016).

Motor

To investigate a brain network linked to motor function, a meta-analysis on finger tapping tasks was performed by Witt et al. (2008). The brain regions consistently activated were the bilateral sensorimotor cortex, basal ganglia, anterior cerebellum, inferior parietal cortex, left ventral premotor cortex and supplementary motor area (Witt, Meyerand, & Laird, 2008).

Reward (Rew)

A meta-analysis on reward-related decision making revealed a network consisting of the bilateral insula, thalamus, brain stem, mid-orbitofrontal cortex, middle frontal gyrus, right nucleus accumbens, left pallidum, left dorsomedial prefrontal cortex, left medial orbitofrontal cortex, right amygdala, supplementary motor area, anterior and posterior cingulate cortex, left inferior parietal lobe, right angular gyrus, left frontal pole and left superior frontal gyrus (Liu, Hairston, Schrier, & Fan, 2011).

Semantic memory (SM)

Semantic memory entails the knowledge we gained from experience (Binder, Desai, Graves, & Conant, 2009). A meta-analysis revealed that the angular and supramarginal gyrus, middle temporal gyrus, posterior inferior temporal gyrus, mid-fusiform gyrus, parahippocampus, dorsomedial, ventromedial and orbital prefrontal cortex, superior, middle and inferior frontal gyrus, posterior cingulate gyrus and ventral precuneus were predominantly activated by studies employing semantic memory tasks (Binder et al., 2009).

Vigilant attention (VigAtt)

Vigilant attention describes the ability to maintain attention on repetitive and unengaging tasks for which not much cognitive effort is needed (Langner & Eickhoff, 2013). The network underlying this cognitive function consists of the anterior paracentral lobe, right medial posterior superior frontal gyrus, dorsal midcingulate cortex, bilateral inferior frontal junction, anterior insula, thalamus, right inferior frontal sulcus, left precentral gyrus, left inferior occipital gyrus, right temporo-parietal junction, right middle occipital gyrus, right inferior parietal lobe and cerebellum (Langner & Eickhoff, 2013).

Working memory (WM)

Working memory describes the ability to encode, maintain and retrieve information over a short period of time. Brain regions commonly activated by working memory tasks include the bilateral anterior insula, inferior frontal gyrus, caudal and rostral lateral prefrontal cortex, posterior superior frontal gyrus, thalamus, cerebellum, intraparietal sulcus, superior parietal lobe, left nucleus caudate and left globus pallidum (Rottschy et al., 2012).

S3 Peak coordinates in MNI space of networks

| network | node number | x | y | z |
|---|---|---|---|---|
| AM | 1 | -1 | -53 | 21 |
| | 2 | -26 | -28 | -17 |

| | | | |
|---|---|---|---|
| 3 | -49 | -61 | 31 |
| 4 | -2 | 51 | -11 |
| 5 | -60 | -9 | -18 |
| 6 | -50 | 27 | -12 |
| 7 | 26 | -33 | -15 |
| 8 | -1 | 20 | 57 |
| 9 | 55 | -58 | 30 |
| 10 | -47 | 9 | 46 |
| 11 | -42 | 53 | 7 |
| 12 | 26 | -14 | -23 |
| 13 | 52 | -5 | -18 |
| 14 | -39 | 13 | -41 |
| 15 | -38 | -82 | 38 |
| 16 | -48 | 29 | 17 |
| 17 | 52 | 31 | -11 |
| 18 | -11 | 62 | 9 |
| 19 | 4 | -8 | 2 |
| 20 | -4 | 39 | 16 |
| 21 | -5 | -34 | 36 |

| | | | | |
|---|---|---|---|---|
| | 22 | -29 | 16 | 51 |
| | 23 | 31 | 1 | -26 |
| CogAC | 1 | 36 | 22 | -4 |
| | 2 | 2 | 16 | 48 |
| | 3 | 48 | 12 | 30 |
| | 4 | 36 | 2 | 54 |
| | 5 | 48 | 30 | 24 |
| | 6 | -38 | -44 | 46 |
| | 7 | -24 | -66 | 48 |
| | 8 | 40 | -46 | 46 |
| | 9 | 60 | -44 | 24 |
| | 10 | 30 | -62 | 52 |
| | 11 | -44 | 10 | 30 |
| | 12 | -34 | 20 | -4 |
| | 13 | -26 | 2 | 52 |
| | 14 | 6 | -18 | -2 |
| | 15 | -40 | -66 | -10 |
| | 16 | 48 | 19 | 6 |
| | 17 | 8 | 29 | 30 |

| | | | | |
|---|---|---|---|---|
| | 18 | -45 | 27 | 30 |
| | 19 | 11 | 7 | 7 |
| eMDN | 1 | -46 | 6 | 30 |
| | 2 | 50 | 12 | 28 |
| | 3 | -32 | 20 | 2 |
| | 4 | 36 | 22 | 0 |
| | 5 | -4 | 14 | 44 |
| | 6 | 6 | 18 | 46 |
| | 7 | -32 | -52 | 46 |
| | 8 | 32 | -58 | 48 |
| | 9 | 44 | 36 | 20 |
| | 10 | -28 | -4 | 52 |
| | 11 | -44 | 32 | 22 |
| | 12 | 32 | 0 | 52 |
| | 13 | -20 | 6 | 4 |
| | 14 | 10 | -12 | 8 |
| | 15 | -46 | -60 | -10 |
| | 16 | 22 | 6 | 4 |
| | 17 | -10 | -16 | 6 |

| | | | | |
|---|---|---|---|---|
| EmoSF | 1 | 4 | 47 | 7 |
| | 2 | 42 | 25 | 3 |
| | 3 | -42 | 25 | 3 |
| | 4 | 48 | 17 | 29 |
| | 5 | -42 | 13 | 27 |
| | 6 | -2 | 8 | 59 |
| | 7 | 20 | -4 | -15 |
| | 8 | -20 | -6 | -15 |
| | 9 | -20 | -33 | -4 |
| | 10 | 14 | -33 | -7 |
| | 11 | 53 | -50 | 4 |
| | 12 | 38 | -55 | -20 |
| | 13 | -40 | -55 | -22 |
| | 14 | 38 | -76 | -16 |
| | 15 | -40 | -78 | -21 |
| | 16 | -4 | 52 | 31 |
| | 17 | 36 | 25 | -3 |
| | 18 | -38 | 25 | -8 |
| | 19 | 2 | 19 | 25 |

| | | | | |
|---|---|---|---|---|
| | 20 | 0 | -15 | 10 |
| | 21 | -2 | -31 | -7 |
| | 22 | -28 | -70 | -14 |
| | 23 | 46 | -68 | -4 |
| | 24 | -48 | -72 | -4 |
| Empathy | 1 | 2 | 56 | 18 |
| | 2 | -8 | 54 | 34 |
| | 3 | 36 | 22 | -8 |
| | 4 | -30 | 20 | 4 |
| | 5 | 50 | 12 | -8 |
| | 6 | 54 | 16 | 20 |
| | 7 | 50 | 30 | 4 |
| | 8 | -44 | 24 | -6 |
| | 9 | -4 | 18 | 50 |
| | 10 | -2 | 28 | 20 |
| | 11 | -4 | 42 | 18 |
| | 12 | -2 | -32 | 28 |
| | 13 | 52 | -58 | 22 |
| | 14 | -56 | -58 | 22 |

| | | | | |
|---|---|---|---|---|
| | 15 | 22 | -2 | -16 |
| | 16 | 54 | -8 | -16 |
| | 17 | 52 | -36 | 2 |
| | 18 | -12 | -4 | 12 |
| | 19 | 6 | -32 | 2 |
| | 20 | 26 | -26 | -12 |
| | 21 | 2 | -20 | -12 |
| | 22 | 14 | 4 | 0 |
| ER | 1 | 48 | 24 | 9 |
| | 2 | 42 | 21 | 45 |
| | 3 | 9 | 30 | 39 |
| | 4 | 0 | -9 | 63 |
| | 5 | -3 | 24 | 30 |
| | 6 | -33 | 3 | 54 |
| | 7 | -36 | 21 | -3 |
| | 8 | -42 | 45 | -6 |
| | 9 | 63 | -51 | 39 |
| | 10 | -42 | -66 | 42 |
| | 11 | -63 | -51 | -21 |

| | | | | |
|---|---|---|---|---|
| | 12 | -51 | -39 | 3 |
| | 13 | 30 | -3 | -15 |
| | 14 | -18 | -3 | -15 |
| eSAD | 1 | 0 | 38 | 10 |
| | 2 | -24 | -10 | -20 |
| | 3 | 24 | -8 | -22 |
| | 4 | -2 | -52 | 26 |
| | 5 | -2 | 32 | -8 |
| | 6 | -46 | -66 | 18 |
| | 7 | 50 | -60 | 18 |
| | 8 | -2 | 52 | 14 |
| | 9 | -6 | 10 | -8 |
| | 10 | 6 | 10 | -8 |
| | 11 | -2 | 50 | -10 |
| | 12 | -54 | -10 | -20 |
| MNS | 1 | -56 | 8 | 28 |
| | 2 | -54 | 6 | 40 |
| | 3 | 58 | 16 | 10 |
| | 4 | 44 | -54 | -20 |

| | | | | |
|---|---|---|---|---|
| | 5 | -38 | -40 | 50 |
| | 6 | 51 | -36 | 50 |
| | 7 | -1 | 16 | 2 |
| | 8 | -54 | -50 | 10 |
| | 9 | -52 | -70 | 6 |
| | 10 | 54 | -64 | 4 |
| | 11 | 30 | -62 | 63 |
| Motor | 1 | -39 | -21 | 54 |
| | 2 | 41 | -16 | 57 |
| | 3 | -3 | -2 | 54 |
| | 4 | -57 | 2 | 32 |
| | 5 | -53 | -24 | 21 |
| | 6 | 45 | -38 | 48 |
| | 7 | -23 | -7 | 1 |
| | 8 | 25 | -8 | 3 |
| | 9 | -22 | -52 | 26 |
| | 10 | 18 | -54 | -22 |
| Rew | 1 | 12 | 10 | -6 |
| | 2 | -10 | 8 | -4 |

| | | | |
|---|---|---|---|
| 3 | 36 | 20 | -6 |
| 4 | -32 | 20 | -4 |
| 5 | 0 | 24 | 40 |
| 6 | 0 | 54 | -8 |
| 7 | 24 | -2 | -16 |
| 8 | 6 | -14 | 8 |
| 9 | -6 | -16 | 8 |
| 10 | 0 | 8 | 48 |
| 11 | 8 | -18 | -10 |
| 12 | -6 | -18 | -10 |
| 13 | 2 | 44 | 20 |
| 14 | -24 | 2 | 52 |
| 15 | -38 | -4 | 6 |
| 16 | 24 | 40 | -14 |
| 17 | -16 | 42 | -14 |
| 18 | 40 | 32 | 32 |
| 19 | -28 | -56 | 48 |
| 20 | 28 | -58 | 50 |
| 21 | 0 | -32 | 32 |

| | | | | |
|---|---|---|---|---|
| | 22 | -36 | 50 | 10 |
| | 23 | -46 | 42 | -4 |
| | 24 | 30 | 4 | 50 |
| | 25 | -22 | 30 | 48 |
| ToM | 1 | 0 | 52 | -12 |
| | 2 | 2 | 58 | 12 |
| | 3 | -8 | 56 | 30 |
| | 4 | 2 | -56 | 30 |
| | 5 | 56 | -50 | 18 |
| | 6 | -48 | -56 | 24 |
| | 7 | 54 | -2 | -20 |
| | 8 | -54 | -2 | -24 |
| | 9 | 52 | -18 | -12 |
| | 10 | -54 | -28 | -4 |
| | 11 | 50 | -34 | 0 |
| | 12 | -58 | -44 | 4 |
| | 13 | 54 | 28 | 6 |
| | 14 | -48 | 30 | -12 |
| | 15 | 48 | -72 | 8 |

| | | | | |
|---|---|---|---|---|
| VigAtt | 1 | -2 | 8 | 50 |
| | 2 | 8 | 32 | 46 |
| | 3 | 0 | 26 | 34 |
| | 4 | 50 | 8 | 32 |
| | 5 | 40 | 22 | -4 |
| | 6 | 46 | 36 | 20 |
| | 7 | -40 | -12 | 60 |
| | 8 | -46 | -68 | -6 |
| | 9 | -48 | 8 | 30 |
| | 10 | 62 | -38 | 17 |
| | 11 | 8 | -12 | 6 |
| | 12 | 32 | -90 | 4 |
| | 13 | -42 | 12 | -2 |
| | 14 | -10 | -14 | 6 |
| | 15 | 6 | -58 | -18 |
| | 16 | 44 | -44 | 46 |
| WM | 1 | -32 | 22 | -2 |
| | 2 | -48 | 10 | 26 |
| | 3 | -46 | 26 | 24 |

| | | | |
|---|---|---|---|
| 4 | -38 | 50 | 10 |
| 5 | 36 | 22 | -6 |
| 6 | 50 | 14 | 24 |
| 7 | 44 | 34 | 32 |
| 8 | 38 | 54 | 6 |
| 9 | 2 | 18 | 48 |
| 10 | -28 | 0 | 56 |
| 11 | 30 | 2 | 56 |
| 12 | -42 | -42 | 46 |
| 13 | -34 | -52 | 48 |
| 14 | -24 | -66 | 54 |
| 15 | 42 | -44 | 44 |
| 16 | 32 | -58 | 48 |
| 17 | 16 | -66 | 56 |
| 18 | -12 | -12 | 12 |
| 19 | -16 | 2 | 14 |
| 20 | -16 | 0 | 2 |
| 21 | 12 | -10 | 10 |
| 22 | -34 | -66 | -20 |

| | | | | |
|---|---|---|---|---|
| | 23 | 32 | -64 | -18 |
| SM | 1 | -46 | -69 | 28 |
| | 2 | -50 | -56 | 31 |
| | 3 | -64 | -44 | -4 |
| | 4 | -47 | -24 | -17 |
| | 5 | -40 | -12 | -30 |
| | 6 | -8 | -57 | 17 |
| | 7 | -20 | 36 | 44 |
| | 8 | -53 | 27 | -4 |
| | 9 | 54 | -59 | 30 |
| | 10 | 43 | -72 | 31 |
| | 11 | -1 | 51 | -7 |
| | 12 | -5 | 56 | 24 |
| | 13 | -31 | -34 | -16 |
| | 14 | -8 | 29 | -10 |
| | 15 | -46 | 25 | 23 |
| | 16 | 64 | -41 | -2 |
| | 17 | -43 | -53 | 55 |
| | 18 | -1 | -18 | 40 |

| | | | |
|---|---|---|---|
| 19 | -2 | -56 | 46 |
| 20 | 51 | 20 | 26 |
| 21 | 64 | -38 | 32 |
| 22 | -23 | 26 | -16 |
| 23 | -5 | -39 | 40 |

S4 Nodes of all networks

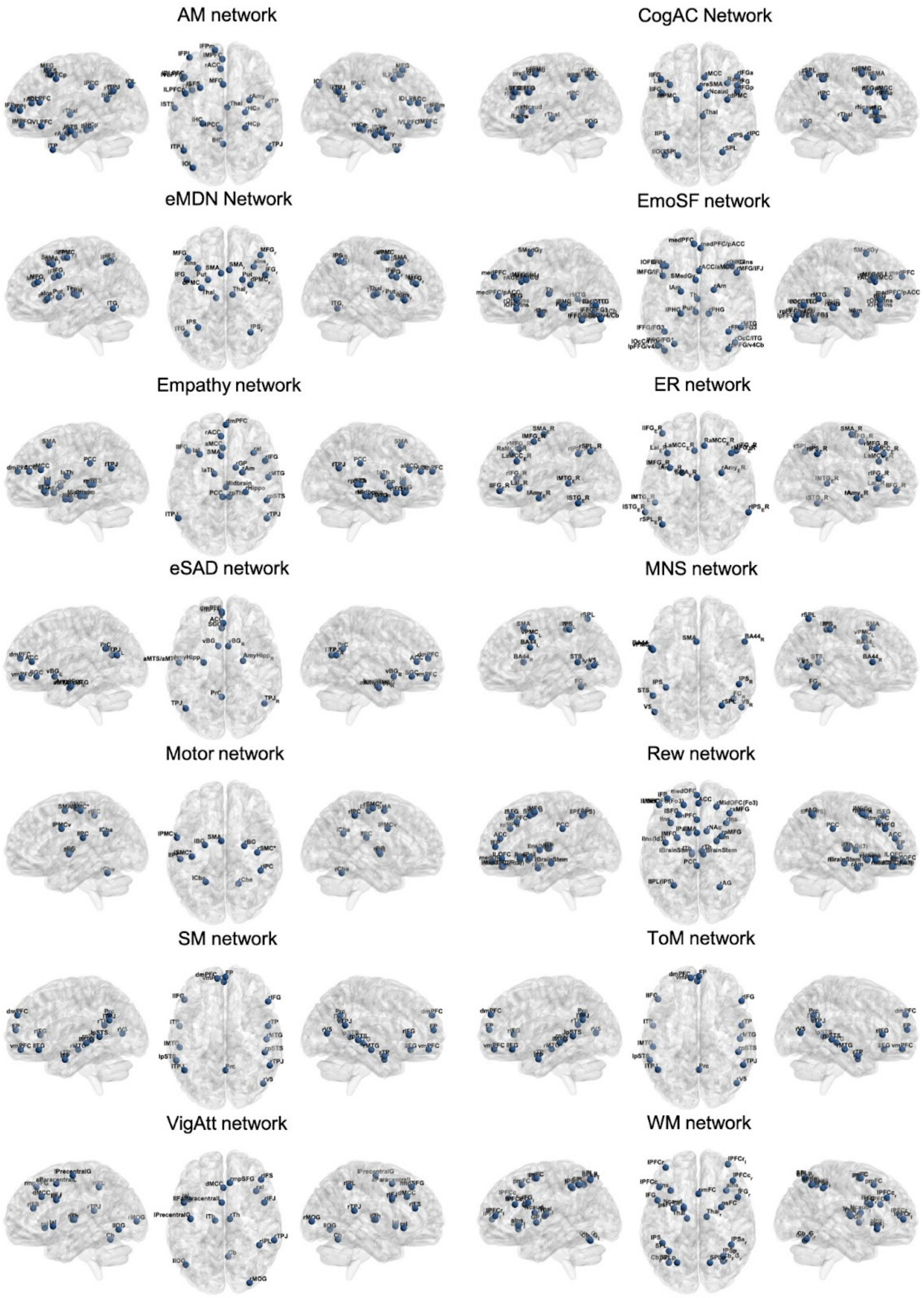


Supp. Figure S4. Visualization of networks nodes of all 14 meta-analytic networks. For peak coordinates of each node, see S3.
